# Privacy-Preserving Pangenome Graphs

**DOI:** 10.64898/2026.02.16.706152

**Authors:** Jacob Blindenbach, Shaunak Soni, Gamze Gürsoy

## Abstract

The human pangenome reference, often represented as a graph, promises to capture genetic diversity across populations, but open release of individual haplotypes raises significant privacy concerns, including risks of re-identification and inference of sensitive traits. To address these challenges, we introduce PanMixer, a framework for privacy-preserving pangenome graph releases that selectively obfuscates an individual’s haplotypes while retaining the utility of the reference graph. PanMixer formulates the privacy–utility trade-off as a knapsack problem, where privacy risk is quantified using information theory and utility is measured using graph properties. Using the recently released draft human pangenome containing 47 individuals, we show that PanMixer robustly reduces re-identification risk under linkage attacks and genome reconstruction attempts. We also show that PanMixer preserves the accuracy of key downstream applications, including allele frequency estimation, linkage disequilibrium analysis, and read mapping. By addressing privacy concerns, PanMixer enables the inclusion of individuals, particularly those from underrepresented populations, who might otherwise be reluctant to contribute but seek representation in future genomic studies. Our results provide both a practical tool and a generalizable framework for balancing privacy and utility in future large-scale pangenome references.

## 1 Introduction

The human pangenome represents a comprehensive reference framework that integrates genomic sequences from diverse populations to capture the full spectrum of human genetic variation [27, 8, 5]. Unlike the current linear reference genome, which is derived predominantly from a limited number of individuals of European ancestry [35], the pangenome incorporates structural variants, insertions, deletions, and population-specific haplotypes that were previously underrepresented or absent [27]. This expanded representation improves read mapping [36], variant calling [7], and functional annotation [19] across ancestries, and consequently enhances the accuracy of genomic analyses. By providing a more inclusive and complete model of human genetic diversity, the pangenome has the potential to advance studies of evolution, disease susceptibility, and precision medicine, and ultimately facilitate equitable applications of genomics in global health [8, 5, 27].

Pangenome graphs are the graphical representation of pangenome references by compressing multiple genomes into nodes of shared DNA sequences connected by edges that capture haplotype specific paths [27]. In 2023, the first human pangenome was released containing 47 individuals[27], along with graph representations generated with two state-of-the-art tools: Pangenome Graph Builder (PGGB) [12] and Minigraph-Cactus [18]. The Human Pangenome Reference Consortium (HPRC) is now expanding this resource to 350 individuals [41], a scale designed to further reduce reference bias and enhance studies of human evolution, disease, and ancestry, with particular relevance for historically underrepresented populations [30]. Public availability of human reference pangenome is critical for accelerating scientific discovery and democratizing data access.

The open release of new individual whole-genome sequence data may raise public concern without some form of privacy protections in place to minimize the risk of re-identifying sample donors. Furthermore, individuals may not be willing to participate in the HPRC under a fully open access model, but these may be critical individuals to include in a pangenome reference resource that is intended to represent all possible haploblocks in the human genome. Such concerns are not merely hypothetical, but are grounded in historical cases where inadequate attention to privacy and consent led to lasting harm. For example, Henrietta Lacks’s cells (HeLa), which were taken and shared without consent, contributed to biomedical breakthroughs such as the polio vaccine [39] and advances in cancer treatment [43, 4], but also created lasting mistrust among African Americans [24, 38]. Similarly, misuse of Native American DNA such as the unauthorized use of Havasupai Tribe samples for research beyond diabetes led to lawsuits, damaged trust, and reduced willingness to participate in genetics studies [6, 2, 13]. To ensure the pangenome represents all populations especially those historically underrepresented, privacy must be front of mind and built in from the start.

In the current pangenome release model, an individual’s genome is represented as haplotype paths through the pangenome graph. Following an individual’s haplotype path through the graph fully reconstructs their genome sequence (Fig. 1A). While this enables scientific utility, it also creates privacy risks: rare or unique variants along a haplotype path can serve as identifiers, enabling re-identification or linkage to sensitive phenotypic traits, as previous studies have shown [9]. Thus, a central challenge emerges: how can individuals contribute to a public pangenome reference in a way that preserves both representation and protects against privacy risks?

**Figure 1.**
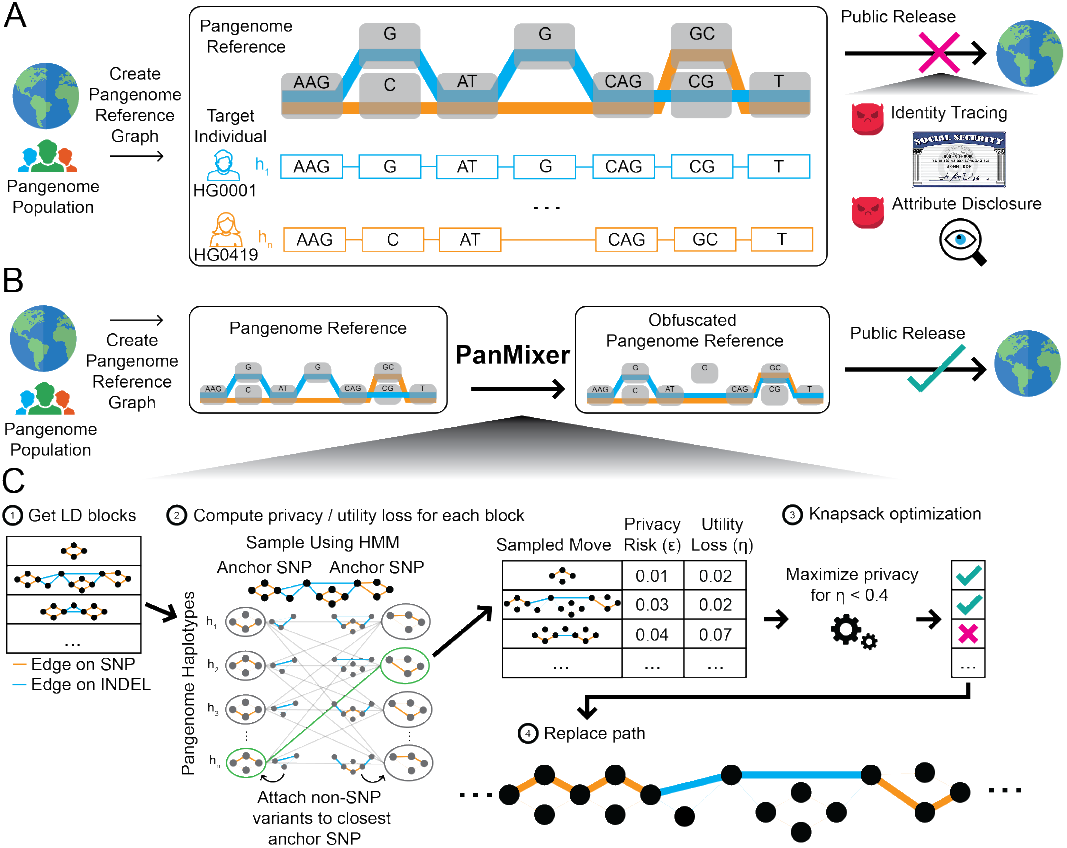
**A:** In the current pangenome release model, a cohort of patients is gathered, sequenced, combined into a pangenome reference using a pangenome graph builder. This pangenome graph where individual haplotypes are represented as paths traversing a sequence of nodes containing varying lengths of DNA is released to the public. Although releasing both the graph and the haplotype paths enables comprehensive representation of population-level variation, an individual’s haplotype paths may reveal private or rare variants that risk re-identification or sensitive attribute inference. **B:** PanMixer updates this release model by obfuscating target individuals paths before release to improve the privacy of these target individuals. **C:** PanMixer (1) identifies LD blocks, (2) computes privacy and utility metrics for potential obfuscation moves sampled via a Li–Stephens-like HMM to preserve LD structure, (3) selects an optimal subset of moves via knapsack optimization given a utility budget, and (4) applies these moves to produce the released obfuscated paths.

To address these privacy concerns, we introduce **PanMixer**, an obfuscation tool that enables participation in pangenome reference while mitigating re-identification risk. PanMixer selectively obfuscates haplotype paths of target individuals to preserve privacy while maintaining the structure and the analytical utility of the graph. We formalized the privacy–utility trade-off as a knapsack problem and, using an integer programming approach, identified the optimal set of paths that minimizes privacy risk while maximizing graph utility for downstream analyses (Fig. 1B). Privacy is quantified using pointwise mutual information, while utility is measured by graph edit distance. We further show that our information-theoretic privacy metric correlates with the success of linkage attacks and our utility metric correlates with the performance of multiple downstream applications, including allele frequency (AF) analysis, linkage disequilibrium (LD) analysis, and read mapping.

## 2 Results

### 2.1 PanMixer is a framework designed to balance genomic privacy and data utility in pangenome graphs

Safeguarding individual privacy while maintaining the utility of genomic resources is a central challenge for the next generation of pangenome projects. PanMixer addresses this challenge by introducing a principled framework for selectively obfuscating haplotype paths within a pangenome graph. The input to PanMixer is the haplotype paths of a target individual whose privacy we want to protect, and the output is a privacy-preserving version of the pangenome graph. PanMixer achieves this by defining a set of obfuscation moves on a target individual’s haplotype paths. These moves modify the nodes and edges the haplotype path of a certain individual traverses inside the graph. Each move improves the privacy of the target individual at a utility cost to the pangenome graph. At its core, PanMixer solves an optimization problem, finding a subset of obfuscation moves that achieves the desired level of privacy for the target individual while minimizing the loss of utility of the pangenome graph. We formulate this optimization as the 0–1 knapsack problem. In the 0-1 knapsack program, someone seeks to choose a subset of items, each with a weight and value, that maximizes the total value without exceeding the knapsack’s capacity [29]. Analogously, each “item” is an obfuscation move, the “weight” corresponds to the utility loss this move causes to the pangenome graph, and the “value” represents the privacy risk to the target individual (Fig. 1C). In PanMixer, we measure utility loss using the Weighted Path Edit Distance (WPED) between the original and obfuscated paths, and privacy risk using the Pointwise Mutual Information (PMI) between them. We define these terms formally in the following sections.

Under the hood, each obfuscation move is defined on an LD block. These blocks are computed from SNPs on the linear reference backbone of the graph (what we will refer to as top level SNPs) [31]. The obfuscation outcome of each block is obtained from either population level allele frequencies or through a Hidden Markov Model (HMM) from Li-Stephens model [26] if there are enough top level SNPs to model the haplotypes in the LD block. Once we determine all the possible moves, we use OR-Tools [32], a 0-1 knapsack linear optimization solver to find the best set of moves to take that achieve a certain level of privacy for the target individual while minimizing the utility loss of the pangenome graph. These moves replaces the target individuals haplotype paths in the original pangenome graph with a haplotype path that protects the private information leakage for the target individual(Fig. 1C).

### 2.2 Quantifying privacy and utility

We quantify privacy risk (denoted as *ϵ*) using Pointwise Mutual Information (PMI) between the original haplotype path (*h*) and the obfuscated haplotype path (*h*^∗^):

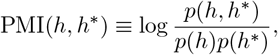

where *p*(*h*) and *p*(*h*^∗^) denote the probabilities of observing the original and obfuscated haplotype paths in the population, respectively, and *p*(*h, h*^∗^) denotes their joint probability. We define the joint probability *p*(*h, h*^∗^) blockwise over independent LD blocks (*b*_1_, · · ·, *b*_*k*_). For blocks that are not obfuscated, the released and original haplotypes are identical, implying the joint probability *of that block* equals the marginal probability 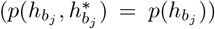. Conversely, for obfuscated blocks, the released haplotype is sampled independently of the original, yielding a factorizable joint distribution 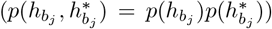. This construction leads directly to the blockwise decomposition of PMI derived in Supplementary Lemma 1. A comprehensive list of all mathematical notations used in this paper is provided in Supplementary Table S1.

High PMI corresponds to high privacy risk as the obfuscated path is strongly statistically dependent on the original path. This makes it easier for an attacker observing the obfuscated path to reconstruct the original path. By contrast, a small PMI or even a PMI of PMI(*h, h*^∗^) = 0 means that the obfuscated path is nearly or is statistically independent of the original and conveys little to nothing about the original path.

For utility (denoted by *η*), we use the Weighted Path Edit Distance (WPED), defined as the minimum cost of all edits required to transform *h* into *h*^∗^. Formally:

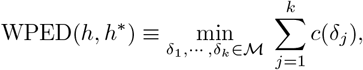

where ℳ defines the set of all possible obfuscation moves, *δ*_*j*_ is an obfuscation move, and *c*(*δ*_*j*_) is its cost. WPED is conceptually similar to the Weighted Graph Edit Distance (WGED), but instead of measuring edits between nodes and edges of two graphs, it measures edits between the original pangenome graph and the graph with obfuscated haplotype paths. A low WPED indicates few edits (high utility), whereas a high WPED indicates many edits (low utility).

#### 2.2.1 Privacy and utility loss in practice

From our equations above, computing privacy risk requires estimating haplotype path probabilities *p*(*h*). Since no large-scale dataset of path frequencies exists, we cannot compute these directly. Instead, we represent haplotypes by the sequence of variants they traverse. In this representation, the graph is decomposed into top level variants and nested variants (variants occurring within the span of larger variants, such as a SNP inside a large insertion). Each haplotype path corresponds to a sequence of alleles across these variants. This variant-centric view allows us to approximate path probabilities from population allele frequencies.

We use the 1000 Genomes dataset for top level SNP frequencies, and the PanGenie callset [7] for structural and nested variants, derived from sequencing reads of all 1000 Genomes individuals aligned to the draft pangenome graph [27]. These AF estimates are then integrated with an HMM that accounts for LD between variants to get an overall probability for each haplotype path.

Privacy risk is therefore tightly linked to allele frequencies, with rare variants contributing disproportionately to the privacy risk and being obfuscated first. This dependency ensures that PanMixer behaves conservatively for individuals from underrepresented populations or for novel variants where external reference data may be sparse. When external frequency estimates are unavailable, PanMixer derives estimates from the graph itself. Thus, an individual with an LD block that contains a novel variant will have high privacy risk for that LD block, and its corresponding obfuscation move will be prioritized for selection.

For utility, we again rely on the variant-centric representation of the graph. Obfuscation moves are restricted to allele modifications within LD blocks (including nested variants). Each move has cost 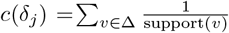, where Δ is the set of variants altered by the move, and support(*v*) is the number of haplotypes traversing variant *v*. This cost function closely approximates AF changes, and for LD blocks with multiple or nested variants, the total utility loss is roughly proportional to the number of edits. Consider a single SNP (Supplementary Fig. S4) on a linear reference backbone such as GRCh38, where an individual’s haplotype follows the reference path. Redirecting it through the alternate allele changes its AF by 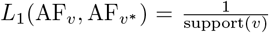, where *L*_1_ is the L1 loss function and *n*_*a*_ is the count of alternate alleles. The resulting utility loss is constant and does not depend on AF, in contrast to privacy risk, which scales with AF.

Because utility loss scales predictably with the number of edits, whereas privacy risk depends strongly on population allele frequencies, balancing the two requires a formal optimization framework. Each obfuscation move *δ*_*j*_ on an LD block bj is associated with a utility cost *η* _*j*_ = *c(δ* _*j*_ ) and a privacy risk 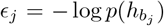(derived in the Supplementary Lemma 2), where 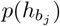 is the population probability of haplotype block 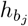. The total privacy risk of the full haplotype is then 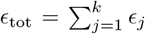. Obfuscating block *b*_*j*_ removes its contribution *ϵ*_*j*_ from the total risk. We therefore define the *removed privacy risk as* 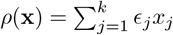, where x ∈ {0, 1}^*k*^ is a binary decision vector indicating which of the k possible obfuscation moves are applied. The realized privacy risk is *ϵ*_res_(x) = *ϵ*_tot_ − *ρ*(x). Minimizing the realized privacy risk is equivalent to maximizing the removed privacy risk. This yields the 0-1 knapsack formulation:

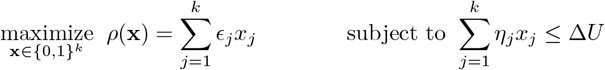

where Δ*U* is the maximum allowable utility loss. The objective is to maximize privacy by removing as much risk as possible, while ensuring that the total utility loss remains within budget.

### 2.3 Symmetric privacy–utility tradeoffs reveal robustness of PanMixer optimization

We generated and characterized the privacy–utility tradeoff in pangenome graph obfuscation by applying PanMixer 44 times for all 44 individuals in the HPRC v1 pangenome reference [27] graph created using PGGB [12]. For each target individual, we defined a set of normalized utility loss levels ranging from 0 (unaltered reference) to 1 (maximally perturbed, where all variants are modified), and optimized for privacy (we have illustrated this process schematically in Fig. 2A). To ensure consistent comparisons across individuals, we similarly normalized the resulting privacy risk to the range [0, 1] by dividing by the total privacy risk of the individual’s original haplotypes (*ϵ*_tot_).

**Figure 2.**
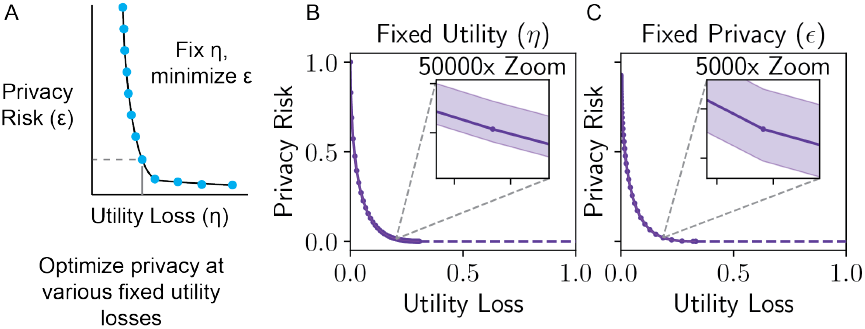
Privacy–utility tradeoff across individuals and ancestries. **A:** Schematic of the privacy–utility tradeoff curve generation process. We picked various utility-loss capacities, and then minimized the privacy risk (*ϵ*) for a target individual constrained by that utility loss (*η*). **B:** Privacy risk (*ϵ*) achieved by PanMixer across a range of fixed utility-loss levels, averaged over all individuals (*n* = 44). Shaded region denotes the full range across individuals; solid line shows the mean. The privacy risk has been normalized to [0, 1] by dividing by the total privacy risk, *ϵ*_tot_ (not having any obfuscation). **C:** Reciprocal optimization: utility loss incurred under fixed privacy constraints. Tradeoff curves are nearly identical, confirming symmetry in the optimization problem and consistency across ancestries.

We observed a consistent concave tradeoff shape, with diminishing returns in privacy as utility loss increased (Fig. 2B). The apparent smoothness of the privacy–utility curves reflects the heterogeneous distribution of block-level privacy contributions rather than a trivial optimization, and no single operating point is privileged by design. The solid line denotes the mean privacy achieved across individuals, and the shaded region captures the full individual-level range. Stratifying by ancestry revealed almost no visible difference between the tradeoff curves (See Supplementary Fig. S1), indicating that our definitions of privacy and utility as well as the optimization generalizes well across all populations.

We confirmed the robustness of the optimization by repeating the analysis under fixed privacy risk, optimizing instead for minimal utility loss (Fig. 2C). The resulting curves closely matched those from the privacy-optimized procedure, demonstrating the symmetry of the tradeoff and validating PanMixer’s consistency across optimizing for privacy and utility.

### 2.4 Privacy metric align with linkage attack and genome reconstruction risk

Next, we show how our privacy metric is indicative of the re-identification risk. To demonstrate this empirically, we evaluated the pangenome graphs produced by PanMixer under an adversarial model in which linkage attacks were attempted, assessing whether obfuscated haplotype paths could be connected to genomes in an external genotype database.

As shown in Fig. 3A, the attacker attempts to link each obfuscated haplotype path to genomes in the 1000 Genomes Project Phase 3 callset [1], using a rarity-weighted matching score that emphasizes shared rare variants, similar to approaches previously used [15, 17, 16, 40]. We applied this attack to pangenome graphs obfuscated by PanMixer at varying privacy levels, ranging from 0 (perfect privacy) to 1 (no obfuscation). For each individual with genotypes in the callset, we extracted shared SNPs between the obfuscated path and the database entries, computed rarity scores, and ranked candidates in the database. Finally, we calculated a Gap Score, defined as the linking score between the original and obfuscated haplotypes minus the highest linking score found in the attack database; a re-identification was considered successful if the Gap Score was positive (*>* 0) and unsuccessful if the score was negative (*<* 0).

**Figure 3.**
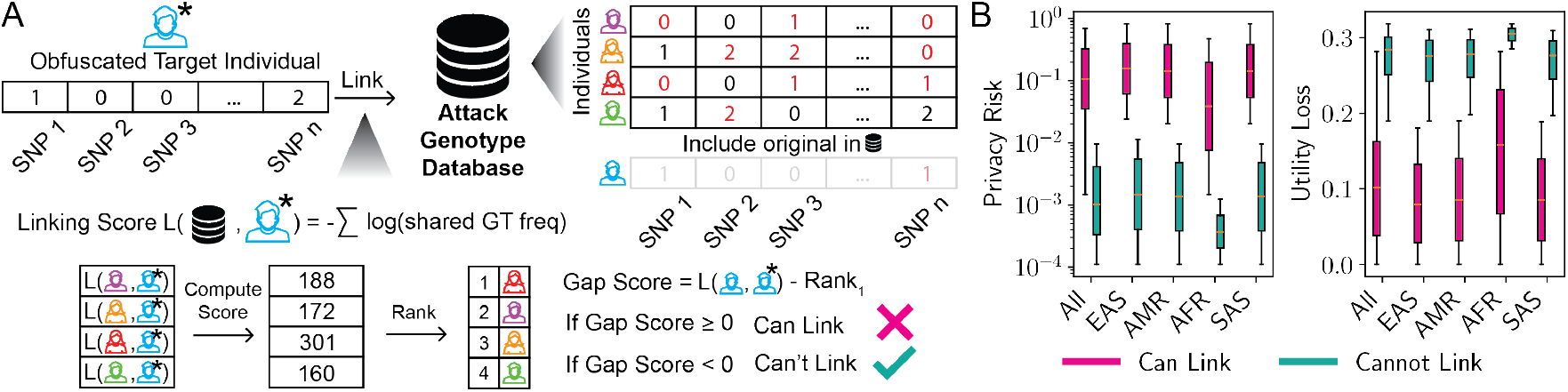
**A:** Schematic of a re-identification attack. An adversary attempts to link an obfuscated individual in the released pangenome graph to an entry in an external genotype database using a rarity-weighted matching score. **B:** Box plots of privacy risk (*ϵ*) and utility loss (*η*) for individuals successfully linked versus not linked by the attacker and utility loss (*η*) for individuals successfully linked versus not. Higher privacy risk (*ϵ*) is associated with increased re-identification success, confirming that individuals who are successfully linked retain higher PMI values (less privacy) than those who are safe. This validates the empirical relevance of *ϵ* as a meaningful privacy metric.

In this pangenome graph, linking attacks failed once the privacy risk was reduced below *ϵ* ≤ *ϵ*_private_ = 0.001 (Fig. 3B). We define *ϵ*_private_ as the smallest privacy risk value at which **all** obfuscated target individuals could no longer be linked. Importantly, *ϵ*_private_ is not universal as it depends on the specific graph, external genotype database, and target individuals. PanMixer provides a systematic way to compute *ϵ*_private_ for any given set of target individuals, pangenome graphs, and external genotype databases. In our experiments, reaching this privacy threshold corresponded to an average utility loss of 0.28 (on a scale where the maximum possible loss is 1).

A different class of threats comes from genome reconstruction attacks, where an adversary tries to reconstruct obfuscated alleles from surrounding LD structure to either recover the original alleles or strengthen re-identification attacks. PanMixer obfuscates entire LD blocks rather than individual variants to minimize the ability of such an attacker. Genotype imputation methods [3, 20, 28, 21] rely on LD to infer missing alleles by exploiting correlations with surrounding variants. By splitting the haplotype paths into blocks with little to no LD between them, PanMixer makes the reconstruction of obfuscated LD blocks ineffective using traditional imputation techniques. We demonstrate this empirically by simulating an attacker attempting to reconstruct obfuscated haplotype paths using Beagle [3] and found their accuracy is no better than simply guessing the major allele at each site, which is essentially equivalent to having no information about the true genotype (Supplementary Fig. S2). Thus, LD based methods cannot recover additional information beyond what is already contained in the obfuscated block, simplifying both the obfuscation problem and our privacy analysis to work on a block by block basis.

To demonstrate the necessity of haplotype-level obfuscation, we compared PanMixer against a baseline strategy that only removes unique or private variants while leaving common variation intact [22]. As shown in Supplementary Fig. S3, this strategy failed to prevent re-identification, maintaining a positive Gap Score comparable to the original graph. This confirms that removing unique markers is insufficient for privacy; the unique combination of common alleles within a haplotype remains a quasi-identifier that PanMixer effectively disrupts.

### 2.5 PanMixer obfuscation preserves utility in common pangenome downstream analysis

Utility, like privacy, must be measured in context. To this end, we assessed the impact of PanMixer’s obfuscations on three key downstream features of pangenome analysis: allele frequency spectra, linkage disequilibrium patterns, and read mapping quality.

#### 2.5.1 Impact of obfuscation on AF spectra and linkage disequilibrium

We evaluated the fidelity of AF and LD in pangenome graphs with obfuscated paths produced by PanMixer. Specifically, we compared the change in AF and LD across all variants between the original pangenome graph and the pangenome graphs in which the genome of a target individual was obfuscated at varying levels of utility loss. We measure the fidelity of AF and LD in pangenome graphs using the following fidelity metric: Wasserstein divergence for the AF spectra difference and the L1 loss (sum of absolute differences) between LD matrices derived from the original and obfuscated graphs. LD was measured as pairwise *R*^2^ correlations between SNPs within sliding 5 kb windows.

For AF spectra, we computed the Wasserstein divergence between the AF (AF) distributions of the original and edited graphs for all variants and variant types. Across various values of privacy risk, the divergence remained below 0.1 (Fig. 4A), indicating minimal impact on allele frequencies. To benchmark utility preservation, we compared PanMixer against a ’Removed’ baseline, where target individuals were entirely excluded from the graph. This baseline represents the maximum possible utility loss for a given individual. This baseline showed substantially higher divergence. For PanMixer edited graphs, at *ϵ*_private_ = 0.001 (the privacy risk threshold where linking attacks failed), the Wasserstein divergence was 0.004 for all alleles, 0.004 for SNPs only, and 0.002 for rare SNPs (population MAF *<* 0.05) which is close to an 6x smaller than the divergence observed when target haplotypes were removed entirely, 0.027. For pairwise LD values, the LD matrix was on average 0.003 off at *ϵ*_*private*_, lower than the 0.017 observed for the complete genome removal baseline.

**Figure 4.**
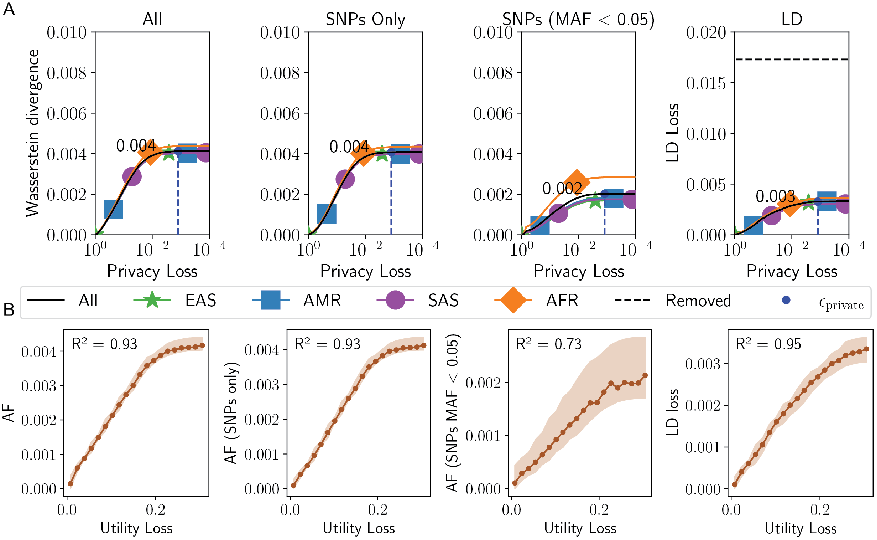
Downstream utility analysis for individuals within the pangenome reference. **A:** Fidelity of allele frequency (AF; all, SNP, and rare SNP variants) and linkage disequilibrium (LD) from obfuscated pangenome graphs across privacy levels (*ϵ*). AF divergence (Wasserstein) remains below0.01 across all *ϵ* but rises sharply under the removed baseline (complete removal of the target’s haplotypes). Likewise, the LD loss stays low (< 0.003) for all targets (*ϵ*_private_). **B:** Correlation between PanMixer’s utility-loss metric and empirical distortions in downstream statistics. Utility loss correlates strongly with both AF divergence (*R*^2^ = 0.93 across all variants; 0.93 for SNPs; 0.73 for rare SNPs) and LD decay (*R*^2^ = 0.95) supporting its validity as an indicator of downstream impact.

#### 2.5.2. Utility loss is correlated with AF and LD divergence

Next, we show the usefulness of our utility metric by showing its correlation with the fidelity metrics (Fig 4B). Utility loss was strongly correlated with AF divergence, with an *R*^2^ of 0.93 for all variants, 0.93 for SNPs only, and 0.73 for rare SNPs (MAF < 0.05). The correlation with LD decay was also strong (*R*^2^ = 0.95).

#### 2.5.3 Obfuscation preserves read mapping quality

We next assessed whether inclusion of an obfuscated genome in the pangenome graph affects its utility for read mapping that originates from sequencing of an external genome. We aligned 1000 Genomes individuals [1] not included in the pangenome graphs to both PanMixer-produced graphs (at *ϵ*_private_, which protects target patients against linking) and to the original graph as a baseline using Giraffe [36] (v1.68.0). Fig. 5A reports the percentage of perfectly aligned reads (all bases matched), gaplessly aligned reads (matched without any indels), and reads with a mapping quality (MAPQ) of 60 [25]. Across all metrics, PanMixer graphs closely matched the baseline with perfectly aligned reads averaged 77.67% (vs. 77.70% with the baseline), gapless reads 95.52% (vs. 95.55%), and reads with a MAPQ of 60 77.36% (vs. 77.29%).

**Figure 5.**
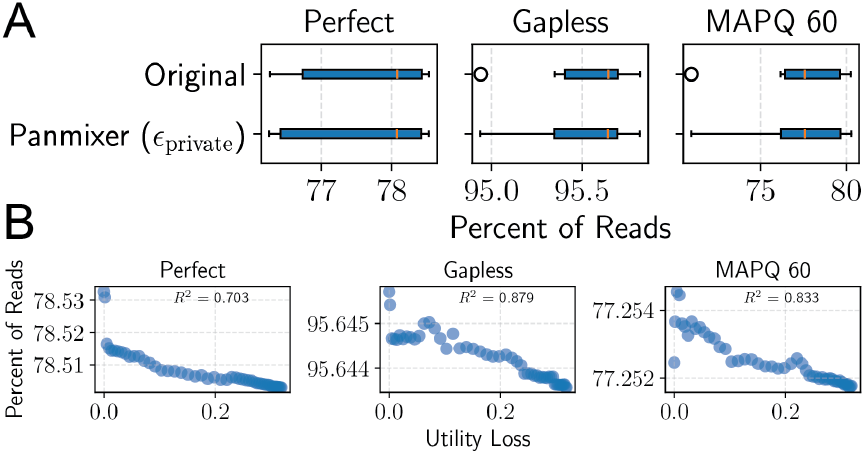
Downstream utility analysis for individuals outside of the pangenome reference. **A:** Percent of reads mapped exactly, gaplessly, or with MAPQ 60 on pangenome graphs generated by PanMixer and the original graph. We aligned 1000 Genomes individuals from five populations (EUR, EAS, AMR, SAS, AFR) against four PanMixer graphs, each obfuscating one target individual (HG00438, HG00733, HG02145, HG03492) at a maximum privacy risk of *ϵ*_private_ = 0.001, preventing linkage to the original. Alignments were performed using vg giraffe [36] (v1.68.0), and mapping statistics were obtained with vg stats. **B:** Correlation between utility loss and percent of perfectly aligned, gapless, and MAPQ 60 reads. Utility loss correlated moderately with perfect alignments (*R*^2^ = 0.703), strongly with gapless alignments (*R*^2^ = 0.879), and strongly with MAPQ 60 alignments (*R*^2^ = 0.833).

We further empirically showed that our utility definition is also correlated with correctly mapped read percentages (Fig. 5B). We found that utility loss correlates strongly with the percent of gapless aligned reads and reads with a MAPQ of 60 (*R*^2^ = 0.879 and *R*^2^ = 0.833 respectively), and moderately with the percent of perfectly aligned reads (*R*^2^ = 0.703).

#### 2.5.4 Scalability of Privacy Preservation

To assess the scalability of PanMixer, we evaluated downstream utility metrics on pangenome graphs with varying numbers of target individuals (*n* = 1 to *n* = 40). We observed that utility loss generally scales linearly with the size of the protected cohort, with notable deviations at higher counts (Supplementary Fig. S6). Specifically, allele frequency divergence (Wasserstein distance) and gapless read alignment exhibited consistent linear degradation. In contrast, both LD loss and the percentage of perfectly aligned reads showed a steady decline that accelerated when nearly every individual was obfuscated (*n* = 40).

High-quality mapping rates (MAPQ 60) increased, suggesting that decreasing graph diversity through obfuscation removes the subtle sequence variations that the target individuals provided. Through the obfuscation of these unique ”features,” the process simultaneously reduces the ability for an attacker to link an individual and trends the graph toward a more linear representation (such as GRCh38) which has been shown to have higher mapping rates than pangenome graphs [37]. Ultimately, these results demonstrate that the cumulative utility cost of privacy is predictable and does not impose a prohibitive burden as the protected cohort size increases.

## 3 Discussion

In this paper, we present PanMixer, the first system to address privacy in pangenome reference releases. PanMixer obfuscates haplotype paths for selected individuals, enabling participation in the pangenome reference without the loss of privacy. We showed that PanMixer can substantially reduce the risk of linking an individual’s genome to external databases under state-of-the-art attacks [15], while preserving the utility of the reference for downstream analyses such as allele frequency estimation, LD analysis, and read mapping.

Unlike exclusion-based strategies or approaches such as removing all unique nodes[22], PanMixer balances privacy and utility by solving a constrained optimization problem. This generates a privacy–utility tradeoff curve that allows informed choices about the level of privacy protection. Our results demonstrate that obfuscated haplotype paths preserve allele frequency spectra, LD decay vectors, and read mapping quality more effectively than simply excluding those unique DNA information.

We also considered Differential Privacy (DP) based approaches, which typically safeguard privacy by injecting statistical noise into the data. While DP is the gold standard for tabular data, applying it to pangenome graphs introduces a critical trade-off between privacy and biological utility. Standard DP mechanisms would require perturbing edges or introducing dummy paths, which breaks the long-range LD structure essential for downstream tasks such as variant calling and read mapping. Recent work has noted that applying DP to pangenome generation often yields sub-optimal results, as the noise required to satisfy privacy bounds destroys the structural integrity of the genome assemblies [22]). Therefore, PanMixer prioritizes the maintenance of topological validity by obfuscating path identifiability through probabilistic switching rather than noise injection, ensuring that the released graph remains biologically useful.

Similarly, while a closed-form analytical expression characterizing the global privacy–utility trade-off would be theoretically appealing, the heterogeneity of genomic data makes such a derivation intractable without imposing unrealistic simplifying assumptions. In PanMixer, privacy risk and utility loss are defined analytically at the local level—specifically, the removed privacy risk for a block is log 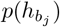 but the global trade-off is governed by the joint distribution of haplotype probabilities and variant supports across the entire genome. Consequently, the relationship between *ϵ* and *η* is determined by a discrete optimization problem over blocks with nonuniform costs and values, which does not admit a simple universal function. Instead, PanMixer provides a precise, data-driven mapping: for any specific pangenome graph, our knapsack formulation computes the exact minimal utility loss required to achieve a given privacy budget *ϵ*. This approach avoids the inaccuracies of loose bounding while capturing the complex, data-dependent structure of real pangenome variation. Future theoretical work could explore deriving analytical bounds under simplified models, such as assuming homogeneous block probabilities or uniform variant supports.

Turning to the practical application of our framework, for curators releasing future pangenome reference graphs, we provide the tools to compute the privacy risk threshold *ϵ*_private_, the point at which linking attacks are no longer successful for any target individual. Importantly, our formulation of privacy risk is data-independent that generalizes across individuals, populations, and pangenome graphs. This enables *ϵ*_private_ to be computed for any pangenome graph and any external database, giving curators a principled and reproducible way to determine when the release of a reference is resilient to re-identification attacks.

It is important to note that the specific privacy threshold (*ϵ*_private_) reported in this study is empirical and dependent on the assumed attacker model, specifically the external reference database (1000 Genomes) and the attack strategy employed. A stronger adversary with access to a larger or more genetically similar reference panel might require a stricter privacy threshold (a lower *ϵ*) to prevent linkage. However, PanMixer is designed not to enforce a universal privacy constant, but to provide a flexible, attacker-agnostic framework. By quantifying privacy via Pointwise Mutual Information (PMI), a metric that captures the fundamental statistical dependence exploitable by linkage or reconstruction attacks, PanMixer allows data owners to simulate various threat scenarios. This shifts privacy assessment from a one-shot empirical claim to a configurable decision process: data owners can explicitly evaluate privacy–utility tradeoffs for the specific threat model they consider relevant to their population and release context, and select an operating point accordingly.

We acknowledge that PanMixer relies on population allele frequency and LD estimates to quantify privacy risk. These inputs may currently be less precise for underrepresented populations or novel variants. However, as pangenome references expand in size and diversity, these estimates will naturally improve. This growth does not impose a computational bottleneck, as PanMixer scales linearly with the number of LD blocks and variants, independent of the total reference cohort size. Future work can build on this framework by evaluating broader adversarial assumptions beyond linkage scenarios, optimizing for additional downstream analyses, and jointly protecting multiple individuals at scale. Further theoretical insight could also be gained by deriving analytical bounds on the privacy–utility trade-off under simplified assumptions.

By addressing privacy directly, PanMixer makes pangenome references more inclusive, secure, and equitable. Lowering the risks of participation can encourage contributions from historically underrepresented populations, broadening the diversity of future genomic resources. In this way, PanMixer not only strengthens the technical foundations of pangenome sharing but also helps ensure that the benefits of genomics are distributed more fairly across all communities.

## 4 Methods

The PanMixer pipeline has three major steps: preprocessing, obfuscation, and optional downstream analysis steps. Preprocessing prepares the graph by splitting it into blocks, obfuscation alters haplotype paths of target individuals using probabilistic models, and downstream analysis evaluates the privacy and utility trade-offs.

### 4.1 Preprocessing

Pangenome graphs are often represented in Graphical Fragment Assembly (GFA) files [14], which specify the DNA sequence of each node, the set of possible edges between nodes, and contig paths as ordered node traversals with orientation. These GFAs are powerful and descriptive of the complex graph substructures that can exist inside of pangenomes supporting inversions, loops, etc. From a privacy perspective, their very lack of structure, the feature that makes them so descriptive, also makes them difficult to safeguard against attacks. We opt to use a more systematic representation such as a Variant Call File (VCF) with nested variants included (produced using tools such as `vg deconstruct` [27]). Tools like PanGenie [7] also use this representations often simplifying further by removing nested variants unless they are on alleles with large reference lengths (*>* 100 kb) [11]. However, we do not remove any variants to preserve some of the complex structures in the graph. We evaluated PanMixer using the Pangenome Graph Builder (PGGB) [12] reference GFA from the initial human pangenome draft [27], running the pipeline on all autosomal chromosomes. Variants were extracted using `vg deconstruct` (v1.36.0) with the `-a` flag to retain all nested variants, using GRCh38 as the backbone reference.

#### 4.1.1 Variant annotations and LD block computation

Let *V* denote all variants and *V*_0_ the set of top level variants (variants on the GRCh38 haplotype path). Variants are categorized as follows:

- *V*_1000G_ ⊂ *V*_0_: Established variants from the Phase 3 phased 1000 Genomes dataset [1], used as trusted anchor points.
- *V*_PG_: Variants identified by variant calling of all 1000 Genomes Phase 3 individuals against the draft human pangenome using the Minigraph-Cactus graph.
- *V*_SNPs_ ⊂ *V*_0_ Subset of variants along the GRCh38 reference backbone that are SNPs.
- *V*_novel_ = *V* \ *V*_SNPs_: Novel variants not represented in existing databases (e.g., 1000 Genomes callset).

Each haplotype block path can be defined using the variants it traverses as we use a variant centric view of the pangenome. LD blocks within the pangenome graph were computed using genotypes from 1000 Genomes Phase 3 variants *V*_1000G_ set. These blocks were computed using `plink` (v.1.9) with the flag `--blocks` to compute LD block boundaries [33, 10]. We grouped variants within the pangenome graph into LD blocks from the boundaries computed by Plink. Variants outside identified LD blocks were treated as individual blocks, and variants not in *V*_1000G_ but within LD the boundaries of an LD block where considered part of that LD block.

### 4.2 Obfuscation

Without loss of generality, we describe the algorithm for obfuscating a single individual inside the pangenome graph. This algorithm can be repeated for any other target individuals inside the pangenome graph. Each individual genome inside the pangenome graph has two haplotype paths which we obfuscate at the same time. For each LD block, we define two obfuscation moves for both haplotype blocks. We budget utility loss by the sum of the obfuscation moves taken on both haplotype paths, and perform all downstream analysis at the genome level.

In summary, for each LD block *j*, we find two obfuscation moves which replace the existing haplotype block path. For each move, (1) we sample a probable new haplotype block, (2) we compute the removed privacy *ϵ*_*j*_, (3) and we compute the utility loss of taking this obfuscation move *η*_*j*_.

#### 4.2.1 Sampling new haplotype block paths

We sample a probable haplotype block path different from the existing haplotype block path of the target individual. This will form a possible move for the obfuscation algorithm to potentially pick. The sampling algorithm is dependent on the number of top level SNPs that exist inside the LD block. If more than 2 exist, enough SNPs exists to build a HMM [26] and we can synthesize more accurate new haplotype blocks. If only one top level SNP exists, we do not have enough useful information to build a HMM. Instead, we sample a new haplotype block path using population allele frequencies.

#### 4.2.2 Sampling using population allele frequencies

For LD blocks containing one or no top level SNPs, we sample alleles directly using population allele frequencies. AF sources depend on the variant type:

- For variants in *V*_1000*G*_, frequencies are taken from the 1000 Genomes dataset.
- For variants in *V*_*PG*_ \ *V*_1000*G*_, frequencies are taken from the PanGenie callset.
- For novel variants not present in either dataset, frequencies are estimated from the pangenome graph itself.

When sampling nested variants, we proceed hierarchically, first sampling from the top level variant, then sampling nested variants conditional on that choice. This maintains structural consistency in the haplotype path.

#### 4.2.3 Sampling a new haplotype block path using an HMM

When at least two top level SNPs are present in an LD block, we construct a Li-Stephens HMM [26] over the donor haplotypes in the pangenome graph (excluding haplotypes belonging to target individuals). The HMM is defined only on top level SNPs *V*_SNPs_, so the resulting sampled haplotype path consists of alleles at those positions (see Fig. 6 for details on the HMM model and parameters used).

**Figure 6.**
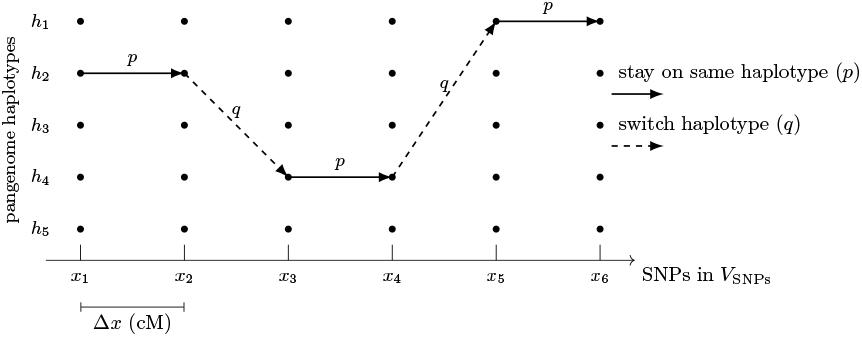
Schematic of the haplotype sampling process using a HMM modeled from Li-Stephens [26] and PanGenie [7]. PanMixer picks a random starting haplotype and probabilistically either stays (solid arrow) or switches (dashed arrow and represents a recombination event). The HMM uses SNPs from *x*_*j*_ *∈ V*_snps_. Model parameters: effective population size *N*_*e*_ = 10, 000 (where *n*_*p*_ is the number of individuals in the pangenome and *n*_*t*_ is the number of target individuals); standard recombination rate *r* = 1.26 [23]; distance between SNPs Δ*x* (centiMorgans); *d* = Δ*x* · *N*_*e*_ · *r*; stay prob-ablity 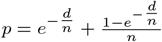switch probability 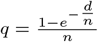 .

To incorporate additional complexity, we attach non-top level variants to the nearest top level SNP. After sampling a donor path with the HMM, we inherit the alleles of these attached variants from the same donor haplotype used at the neighboring to top level SNP.

The final sampled haplotype block therefore combines top level SNPs chosen via the HMM with attached nested or complex variants propagated from the corresponding donor haplotypes yielding a structurally consistent haplotype path.

#### 4.2.4 Computing privacy risk

While PanMixer could directly minimize privacy risk, it instead formulates the optimization problem in terms of maximizing *removed privacy risk*, which is mathematically equivalent. This choice is purely a matter of convention as optimization frameworks such as the knapsack problem are more naturally expressed as maximization problems.

For an LD block *b*_*j*_, the removed privacy risk is given directly by Supplementary Lemma 2 as 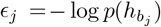, where 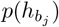 is the population probability of the haplotype path 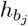.

##### Single top level SNP (or none)

If the LD block contains at most one top level SNP, we compute 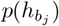 as the product of allele frequencies across all variants *v* in the block: 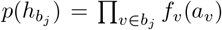, where *a*_*v*_ is the allele of haplotype 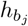 at variant *v* and *f*_*v*_(·) denotes its population frequency. Allele frequencies are obtained as follows:

- For variants in *V*_1000*G*_, frequencies are taken from the 1000 Genomes dataset.
- For variants in *V*_*PG*_ \ *V*_1000*G*_, frequencies are taken from the PanGenie callset.
- For novel variants not present in either dataset, frequencies are estimated directly from the pangenome graph.

For nested variants, the probability is approximated as the product of the population frequency of the parent allele and the marginal frequency of the nested allele. Formally, for a nested variant *v*_nested_ within a parent variant *v*_parent_, we write 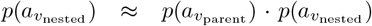, which corresponds to assuming conditional independence of the nested allele given its parent. In practice, this means that the contribution of a nested allele is only counted when its parent allele is present, but the nested AF itself is taken from population level estimates (1000 Genomes, PanGenie, or the graph). While this simplification may not capture true correlations between nested variants and their parents, it provides an approximation in the absence of large-scale conditional frequency data.

##### Multiple top level SNPs

If the LD block contains more than one top level SNP, we can incorporate LD information in computing 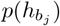. In this case, we compute 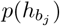 using the Li-Stephens HMM that we use for the sampling process. The haplotype probability can be evaluated with the forward algorithm which estimates the total probability of the observed haplotype path 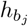 using the top level SNPs in *b*_*j*_ [34].

#### 4.2.5 Computing utility loss

The utility loss of an obfuscation move on a block *b*_*j*_ is 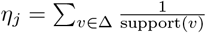where Δ is the set of variants altered by the move, and support(*v*) is the number of haplotypes traversing variant *v*. The utility loss treats nested and regular variants the same as the support only counts the number of present haplotypes ignoring missing and skipped haplotypes.

#### 4.2.6 Obfuscating target individual with knapsack optimization

For all *k* LD blocks, from the process above we have 2 obfuscation moves for each LD block and the reduced privacy risk *ϵ*_*j*_ and utility loss *η*_*j*_ for each move.

We define a binary decision vector on whether to take the move or not as **x** ∈ {0, 1} ^2*k*^. Our realized removed privacy risk and utility loss is a dot product **x** · *ϵ* and **x** · *η*.

We then fix the utility loss as Δ*U*, and solve:

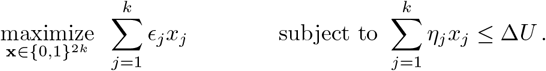

In practice, we utilize the linear optimization solver from OR-Tools [32] and relax the binary constraint on **x**, rounding to 0 or 1 after the computation has completed. All results produced in this paper were from optimal solutions as reported by OR-Tools. The optimization happened on a per chromosome level for efficiency purposes.

### 4.3 Optional downstream Analysis

To evaluate the privacy and utility of the obfuscated pangenome graphs, we performed several downstream utility analysis detailed below.

#### 4.3.1 Relationship between privacy risk and linkage attacks

We evaluated whether target individuals could be empirically linked to external genomic databases (with potentially sensitive attributes) using a privacy attack similar to that described in [15, 17, 16, 40]. Specifically, we attempted to link target individuals to entries in the 1000 Genomes dataset by computing a *genotype linkage score L* between each individual *g*_*i*_ in the 1000 Genomes dataset and a target individual 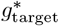.

Although both the 1000 Genomes dataset and the pangenome data are phased, we found that matching at the *genotype* level was the most effective attack strategy, outperforming haplotype level matching (Supplementary Fig. S5).

The genotype linkage score is defined as:

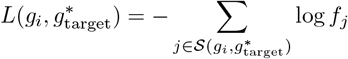

where: 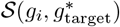 is the set of loci where the genotypes of *g*_*i*_ and 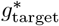 are identical, and *f*_*j*_ is the population frequency of the genotype at locus *j*.

This scoring scheme emphasizes rare allele matches, which are more informative for uniquely identifying individuals. An individual is considered successfully linked if the linkage score between the obfuscated genome and the original genome is greater than or equal to all scores with other database entries:

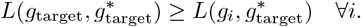

We define the gap score as the difference between the linkage score between the obfuscated genome and the original genome and the linkage score between the obfuscated genome and all other database entries:

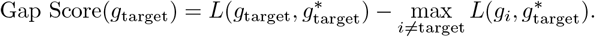

Finally, we determine *ϵ*_private_, the privacy risk threshold at which all target individuals can no longer be linked, by systematically obfuscating each individual across decreasing *ϵ* values until the gap score falls below zero for every individual in the pangenome.

### 4.4 Reconstruction attacks

We simulated a reconstruction attack by modeling an adversary attempting to recover the original haplotype paths from the obfuscated haplotypes. The attacker was assumed to have access to the 1000 Genomes dataset as a reference panel.

For target individuals within our pangenome graph, we first extracted SNPs overlapping with those in the 1000 Genomes dataset. Using only the genotypes at these overlapping SNPs, we ran `beagle` (v5.5) [3] with default parameters for genotype refinement, with the 1000 Genomes dataset (excluding the target individuals) as the reference panel. We then compared the refined genotypes both to the original genotypes and to a baseline reconstruction obtained by selecting the most frequent alleles from the 1000 Genomes dataset.

#### 4.4.1 Relationship between utility loss and allele frequencies

We model each allele at every variant in the pangenome graph as a Bernoulli random variable, and measured the changes in AFs using the Wasserstein divergence between the original and obfuscated allele distributions. For alleles modeled as a Bernoulli random variable, the Wasserstein divergence is two times the mean absolute difference between AFs before and after obfuscation. The factor of two arises because each diploid individual carries two alleles per variant. This approach follows the method of [42], but extends it from the minor allele to all alleles.

Formally, let *p*_*j*_ and *q*_*j*_ denote the allele frequencies at site *j* in the original and graphs with obfuscated individuals, respectively. The AF divergence is defined as: 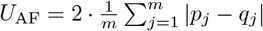, where *m* is the total number of variant sites.

#### 4.4.2 Relationship between utility loss and linkage disequilibrium

We assess changes in LD structure by comparing the LD matrices of the pangenome graphs with obfuscated target individuals to those of the original graph. LD matrices are computed in sliding windows of 5 kb. Let **d**original and **d**obf denote the LD matrices for the original and obfuscated graphs, respectively. The change in LD is defined as the L1 norm between these two matrices: *U*_LD_ = ∥**d**_obf_ − **d**_original_∥_1_ .

#### 4.4.3 Relationship between utility loss and read mapping

We evaluated read mapping performance by aligning sequencing reads from randomly selected 1000 Genomes individuals [1] to pangenome graphs produced by PanMixer. For each population, we randomly selected one individual who was not included in the construction of the pangenome graph: HG00138 (European), HG00635 (East Asian), HG01112 (American), HG02698 (South Asian), and NA18853 (African).

Chromosome 21 reads for these individuals were extracted from CRAM files obtained from the 1000 Genomes repository. The extracted reads were then aligned with Giraffe [36] to the corresponding chromosome 21 graphs to four pangenome graphs each with an target individual from a unique super population obfuscated till linking was no longer possible. The GAM files produced by Giraffe were analyzed using `vg stats` (v1.36.0) to find the percent reads aligned perfectly, gapless, and with a MAPQ of 60.

## 5 Code Availability

The code for PanMixer is available at https://github.com/g2lab/panmixer.

## Supplementary Materials

This document contains the supplementary materials for the paper *Privacy-Preserving Pangenome Graphs*.

### Supplementary Figures

**Figure S1:**
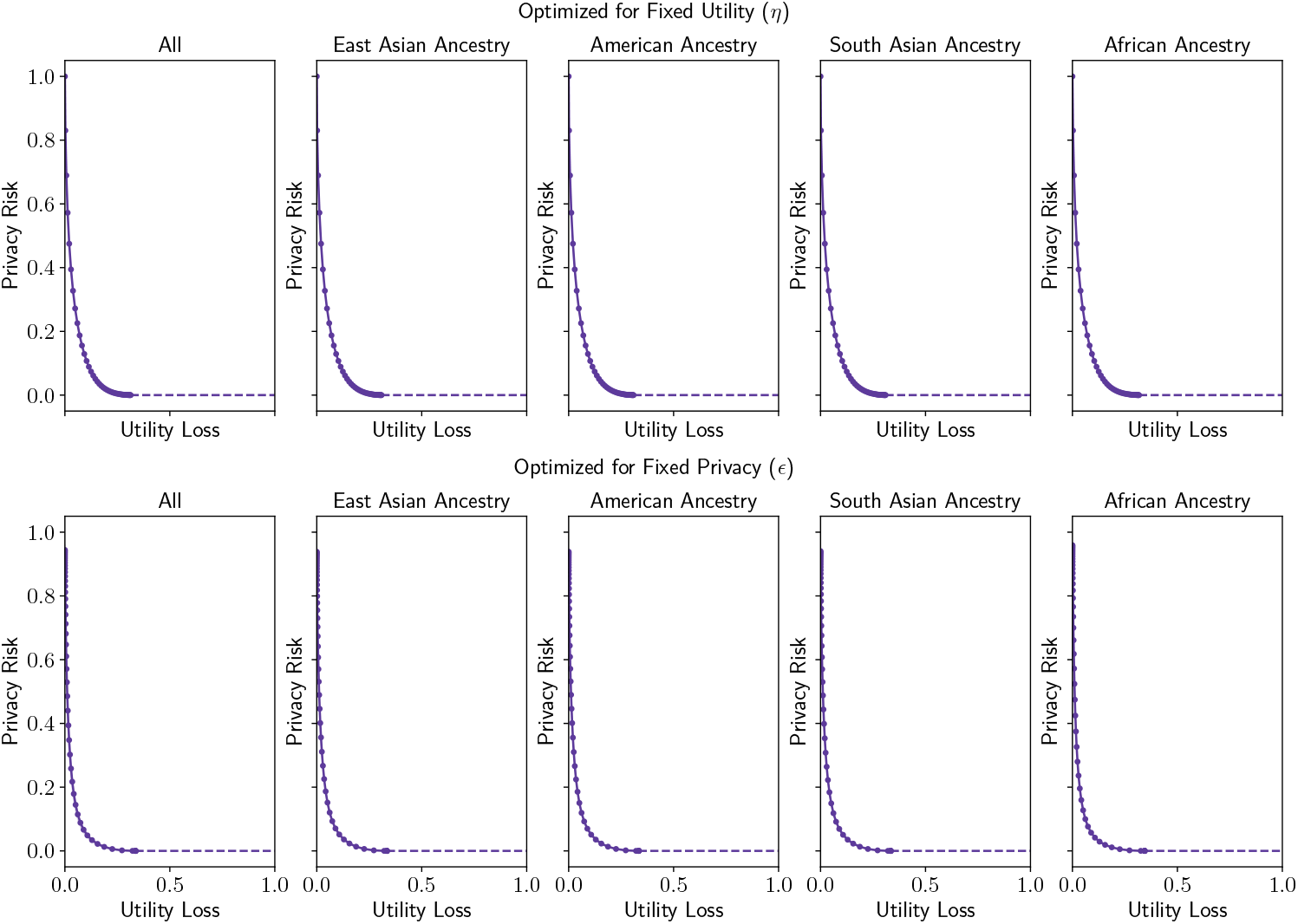
Privacy–utility trade-off across ancestries. Privacy risk (*ϵ*) achieved by PanMixer across a range of fixed utility-loss levels, and utility loss (*η*) achieved across a range of fixed privacy-risk levels. Each plot shows results averaged over all individuals (*n* = 44) and stratified by ancestry: East Asian (*n* = 4), American (*n* = 16), South Asian (*n* = 1), and African (*n* = 23).

**Figure S2:**
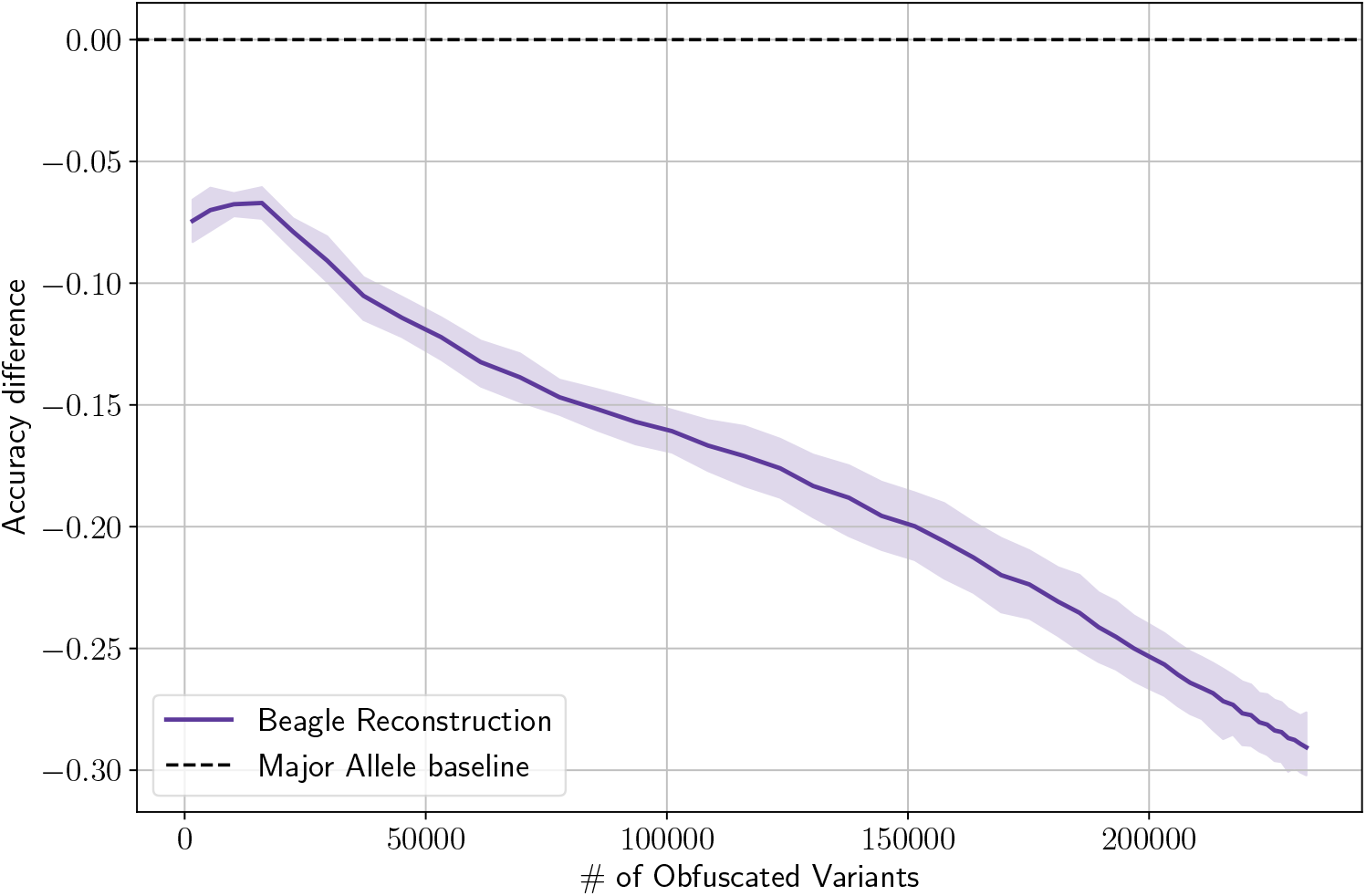
Difference in imputation accuracy between Beagle and a simple major allele baseline across increasing numbers of obfuscated variants. For various privacy loss levels, we compute the mean accuracy difference (Beagle vs. Major Allele). Negative values indicate that Beagle reconstruction performs worse than simply predicting the major allele. Shaded regions highlight uncertainty, and the dashed horizontal line marks parity with the baseline. Results are shown for chromosome 21 reconstructed using Beagle (version 5.5) with the 1000 Genomes genotypes as the reference panel.

**Figure S3:**
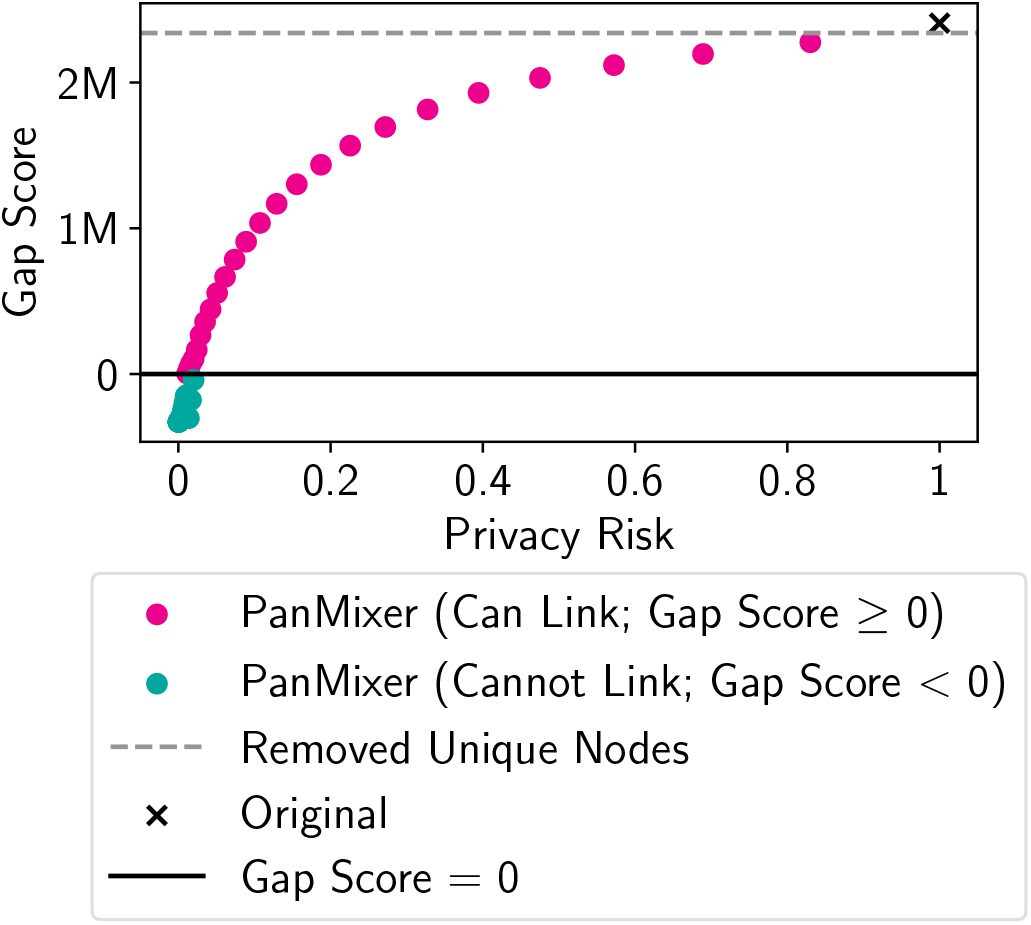
Comparison of re-identification risk across obfuscation strategies. A Gap Score *>* 0 (above the solid black line) indicates successful re-identification, while a score < 0 implies the attack failed. The ”Removed Unique Nodes” baseline (grey dashed line) maintains a high positive Gap Score nearly identical to the Original unobfuscated graph (black cross), demonstrating that removing rare variation alone provides negligible protection against linkage attacks. In contrast, PanMixer (circles) systematically reduces gap scores as the privacy risk decreases. The transition from pink points (can link) to teal points (cannot link) illustrates that PanMixer can successfully drive the Gap Score below zero, rendering the individual unlinkable.

**Figure S4:**
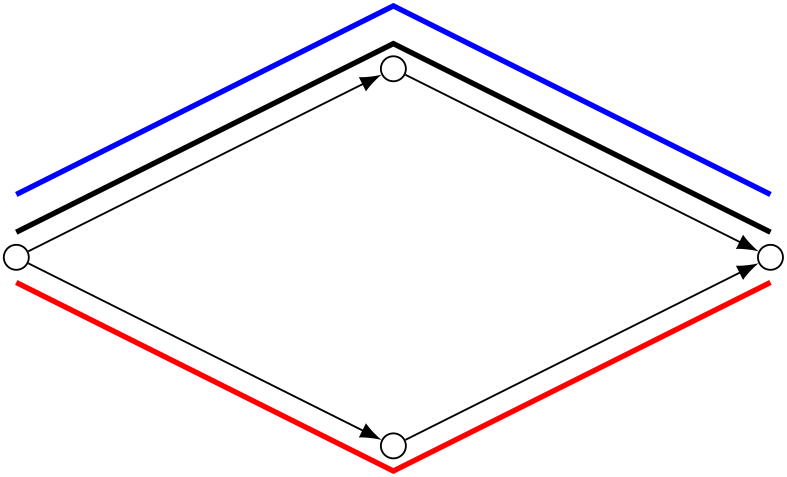
A SNP inside the pangenome graph. The black line represents the linear haplotype reference backbone (such as GRCh38), the blue line represents the target individual’s haplotype path, and the red line represents the obfuscated haplotype path.

**Figure S5:**
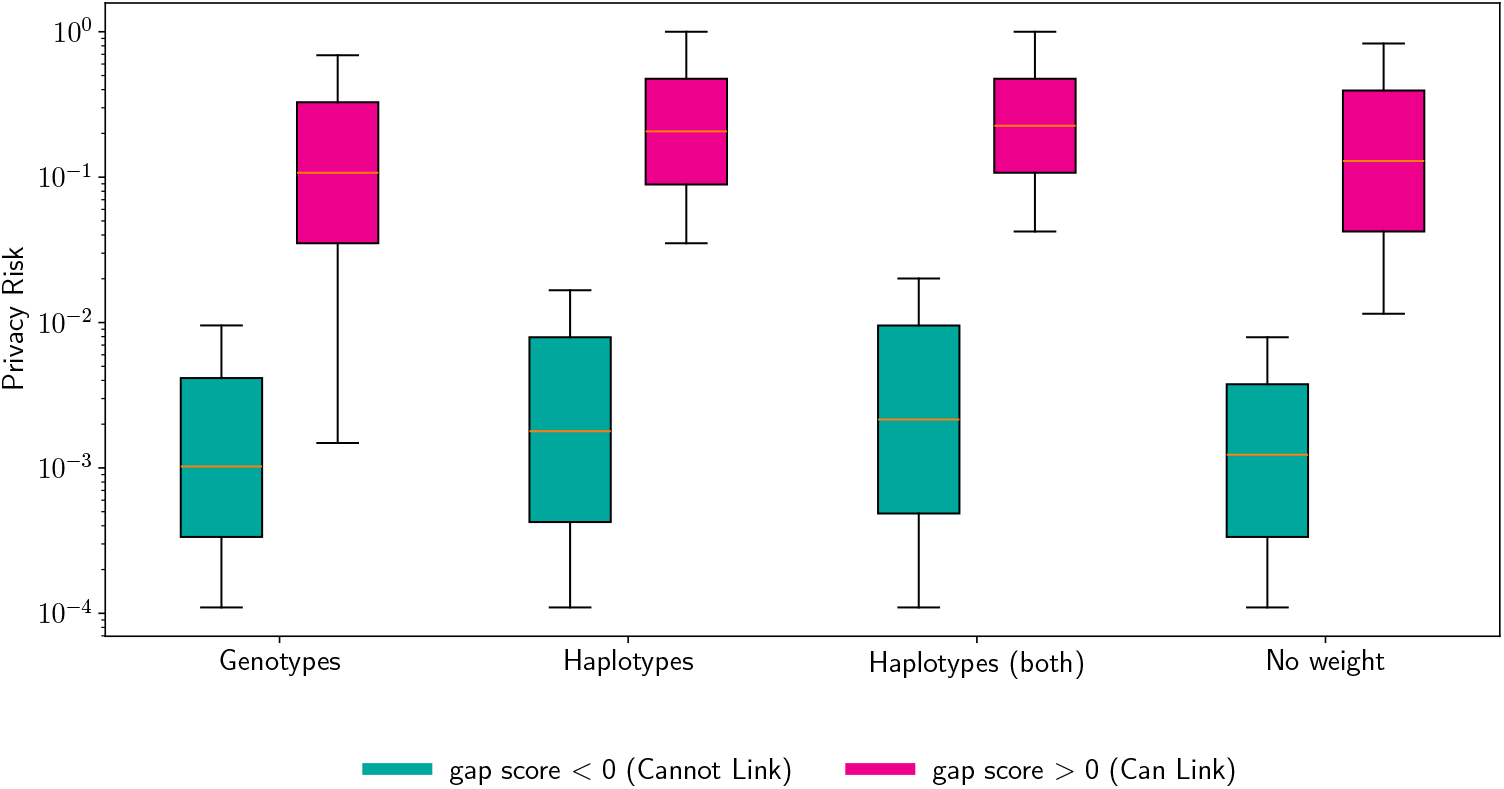
Box plots of privacy risk across all populations for four scoring metrics: *genotypes*, which matches genotypes on the target individual’s genome to those in the attack database; *haplotypes*, which performs this matching at the haplotype level; *haplotypes (both)*, which considers only sites where both haplotypes are equal and phased identically; and *no weight*, which does not weight shared variants by their rarity. For each metric, two distributions are shown: samples with `gap score` < 0 in cyan (cannot link) and `gap score` *>* 0 in purple (can link). Genotype based linking proved most effective, requiring the lowest privacy risk to ensure that the target individual could not be linked.

**Figure S6:**
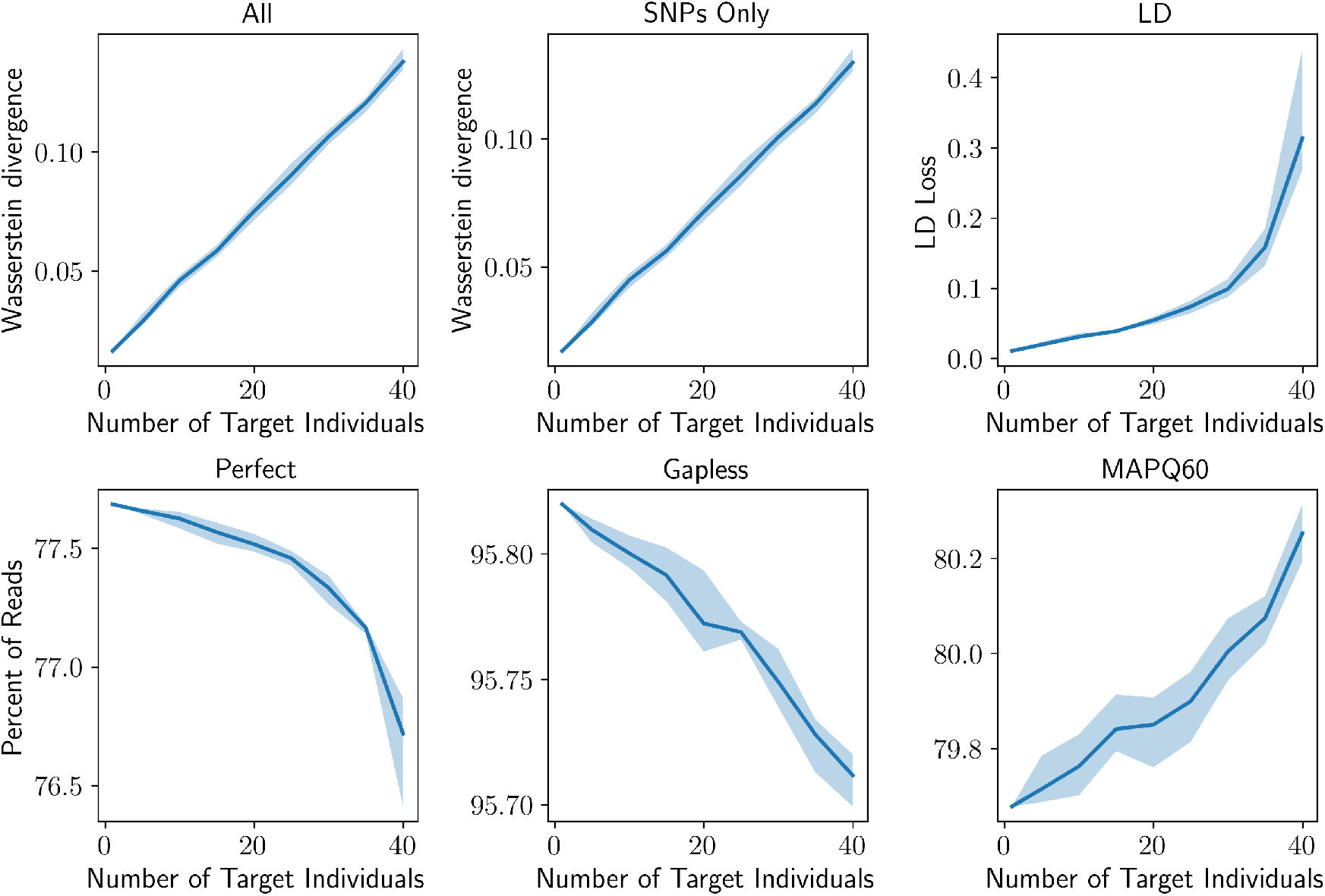
Downstream utility metrics as a function of the number of protected individuals. For each count, we measured metrics across 5 random subsets; the solid line represents the mean and the shaded region indicates the min-max range. The figure displays the impact of simultaneously obfuscating up to 40 target individuals (x-axis) on various genomic utility metrics. Top row: Allele frequency divergence (Wasserstein distance) for all variants and SNPs only, and Linkage Disequilibrium (LD) loss. Bottom row: Read mapping performance, including the percentage of perfectly aligned reads, gapless alignments, and reads with high mapping quality (MAPQ60).

### Supplementary Methods

#### Tables

**Table S1.** Table of Notations.

| Symbol | Description |
| --- | --- |
| *Optimization & Trade-off Variables* |  |
| $\epsilon / \epsilon_{tot}$ | Privacy risk, quantified using Pointwise Mutual Information (PMI) |
| $\eta$ | Utility loss, measured by Weighted Path Edit Distance (WPED) |
| $x$ | Binary decision vector ($x \in \{0, 1\}^{2k}$) indicating selected obfuscation moves |
| $\rho(x)$ | Removed privacy risk, defined as $\sum \epsilon_j x_j$ |
| $\epsilon_{res}(x)$ | Realized privacy risk after obfuscation ($\epsilon_{tot} - \rho(x)$) |
| $\Delta U$ | Maximum allowable utility loss (Knapsack capacity) |
| $\epsilon_{private}$ | Privacy risk threshold where linking attacks fail for all targets |
| *Graph & Haplotype Definitions* |  |
| $h$ | The original haplotype path |
| $h^*$ | The obfuscated haplotype path |
| $b_j$ | The $j$-th Linkage Disequilibrium (LD) block |
| $\delta_j$ | An obfuscation move applied to block $b_j$ |
| $c(\delta_j)$ | The cost associated with obfuscation move $\delta_j$ |
| $\eta_j$ | The utility loss associated with obfuscation move $\delta_j$ |
| $\epsilon_j$ | Privacy risk associated with obfuscation move $\delta_j$ |
| $V$ | Set of all variants in the pangenome graph |
| $V_0$ | Set of top-level variants on the linear reference backbone |
| $V_{SNPs}$ | Subset of $V_0$ consisting of Single Nucleotide Polymorphisms |
| $V_{1000G}$ | Variants established from the 1000 Genomes Phase 3 dataset |
| $support(v)$ | Number of haplotypes traversing variant $v$ |
| *Metrics & Scoring* |  |
| $PMI(h, h^*)$ | Pointwise Mutual Information between original and obfuscated paths |
| $WPED(h, h^*)$ | Weighted Path Edit Distance between original and obfuscated paths |
| $L(g, g^*)$ | Linkage score between a genome $g$ and obfuscated genome $g^*$ |
| $U_{AF}$ | Allele Frequency divergence (Wasserstein divergence) |
| $U_{LD}$ | Change in LD structure ($L_1$ norm of LD matrix difference) |
| *HMM Parameters (Li-Stephens Model)* |  |
| $N_e$ | Effective population size |
| $r$ | Recombination rate |
| $d$ | Distance between SNPs in centiMorgans |
| $p$ | Probability of staying on the same haplotype |
| $q$ | Probability of switching haplotypes (recombination) |

##### Proofs

**Remark 1** (Utility loss is similar to AF). Consider a single SNP (See Supplementary Fig. S4 for an example) inside the pangenome graph on a linear reference backbone such as GRCh38, and the haplotype of an individual taking this reference path. Redirecting the haplotype through the alternate allele changes its AF by

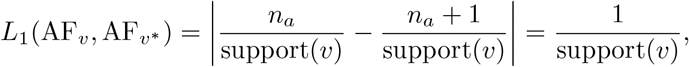

where *L*_1_ is the 𝓁_1_ loss and *n*_*a*_ is the count of alternate alleles. The utility loss is constant for this move and does not depend on AF.

###### Lemma 1

(Blockwise additivity of PMI). Let *h* and *h*^∗^ be haplotype paths decomposed into *k* LD blocks 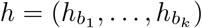 and 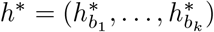. Assume the following blockwise factorizations hold:

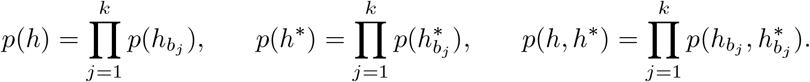

Then the pointwise mutual information (PMI) decomposes as

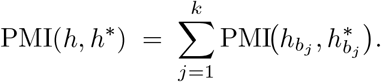

*Proof*. By definition,

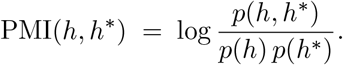

Using the stated factorizations,

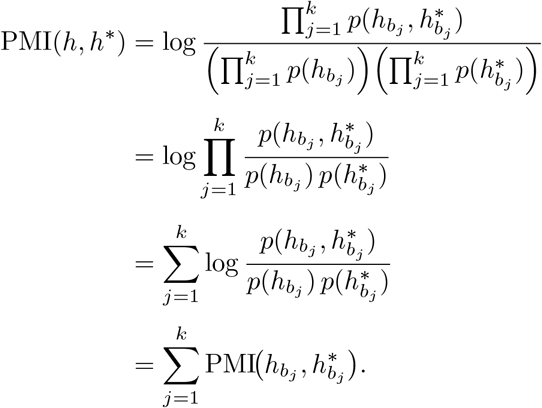

###### Lemma 2

(Removed privacy risk of an obfuscated block). Let *h* be a haplotype path with LD block decomposition 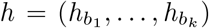, and let *h*^∗^ be an obfuscated version of *h* that differs only at block *j*. Then the removed privacy risk—the reduction in PMI from the original haplotype *h* to its obfuscated counterpart *h*^∗^—is equal to the self-information of the original block:

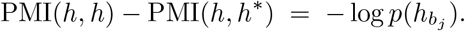

*Proof*. By definition, the removed privacy risk is

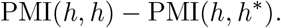

Applying Lemma 1, we expand both terms blockwise:

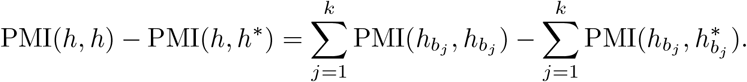

Since *h* and *h*^∗^ differ only at block *j*, all terms cancel except those at *i*:

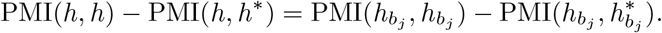

Because the obfuscated block 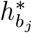 is sampled independently of 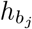, we have 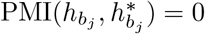 Thus

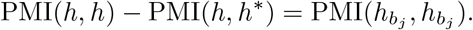

Finally, expanding the definition,

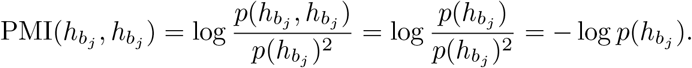

